# Protein sequence design by explicit energy landscape optimization

**DOI:** 10.1101/2020.07.23.218917

**Authors:** Christoffer Norn, Basile I. M. Wicky, David Juergens, Sirui Liu, David Kim, Brian Koepnick, Ivan Anishchenko, Foldit Players, David Baker, Sergey Ovchinnikov

**Author notes:** A list of participants will appear in the supplementary information (Foldit Players). These authors contributed equally (Christoffer Norn, Basile I. M. Wicky).

## Abstract

The protein design problem is to identify an amino acid sequence which folds to a desired structure. Given Anfinsen’s thermodynamic hypothesis of folding, this can be recast as finding an amino acid sequence for which the lowest energy conformation is that structure. As this calculation involves not only all possible amino acid sequences but also all possible structures, most current approaches focus instead on the more tractable problem of finding the lowest energy amino acid sequence for the desired structure, often checking by protein structure prediction in a second step that the desired structure is indeed the lowest energy conformation for the designed sequence, and discarding the in many cases large fraction of designed sequences for which this is not the case. Here we show that by backpropagating gradients through the trRosetta structure prediction network from the desired structure to the input amino acid sequence, we can directly optimize over all possible amino acid sequences and all possible structures, and in one calculation explicitly design amino acid sequences predicted to fold into the desired structure and not any other. We find that trRosetta calculations, which consider the full conformational landscape, can be more effective than Rosetta single point energy estimations in predicting folding and stability of de novo designed proteins. We compare sequence design by landscape optimization to the standard fixed backbone sequence design methodology in Rosetta, and show that the results of the former, but not the latter, are sensitive to the presence of competing low-lying states. We show further that more funneled energy landscapes can be designed by combining the strengths of the two approaches: the low resolution trRosetta model serves to disfavor alternative states, and the high resolution Rosetta model, to create a deep energy minimum at the design target structure.

**Significance:** Computational protein design has primarily focused on finding sequences which have very low energy in the target designed structure. However, what is most relevant during folding is not the absolute energy of the folded state, but the energy difference between the folded state and the lowest lying alternative states. We describe a deep learning approach which captures the entire folding landscape, and show that it can enhance current protein design methods.

## Introduction

Computational design of sequences to fold into a specific protein backbone structure is typically carried out by searching for the lowest energy sequence for the desired structure. In Rosetta and related approaches, side-chain rotamer conformations are built for all amino acids at all positions in the structure, the interaction energies of all pairs of rotamers at all pairs of positions are computed, and combinatorial optimization (in Rosetta, Monte Carlo simulated annealing) of amino acid identity and conformation at all positions is carried out to identify very low energy solutions. Over the past 25 years, a large number of algorithms (1–3) have been developed to solve this problem, including recent deep learning based solutions (4–6). A limitation of all of these approaches, however, is that while they generate a sequence that is the lowest energy sequence for the desired structure, they can result in rough energy landscapes that hamper folding (7, 8), and do not guarantee that the desired structure is the lowest energy structure for the sequence. Thus, an additional step is usually needed to assess the energy landscape, and determine if the lowest energy conformation for a designed sequence is the desired structure; the designed sequences are subjected to large scale stochastic folding calculations, searching over possible structures with the sequence held fixed (9). This two step procedure has the disadvantage that the structure prediction calculations are very slow, requiring for adequate sampling of protein conformational space many CPU-days. Moreover, there is no immediate recipe for updating the designed sequence based on the prediction results—instead sequences which do not have the designed structure as their lowest energy state are typically discarded. Multistate design (10–12) can be carried out to maximize the energy gap between the desired conformation and other specified conformations, but the latter must be known in advance and be relatively few in number for such calculations to be tractable.

We recently described a convolutional neural network called trRosetta that predicts residue-residue distances and orientations from input sets of aligned sequences. Combining these predictions with Rosetta energy minimization yielded excellent predictions of structures in benchmark cases, and in more recent blind CAMEO structure modeling evaluations (13). While co-evolution between pairs of positions in the input multiple sequence alignments is critical for accurate prediction of structures of naturally occurring proteins, we found that it is not necessary for more “ideal” designed proteins: accurate predictions of the latter can be obtained from single sequences (13). Because distance and orientation predictions are probabilistic, we rationalised that they might inherently contain information about alternative conformations, and thus provide more information about design success than classical energy calculations. Moreover, because these predictions can be obtained rapidly for an input sequence on a single GPU, we reasoned that it should be possible to use the network to directly design sequences that fold into a desired structure by maximizing the probability of the observed residue-residue distances and orientations versus all others. Unlike the standard energy-based sequence design approaches described in the previous paragraph, such an approach would have the advantage of explicitly maximizing the probability of the target structure relative to all others (Fig. 1A).

**Fig. 1.**
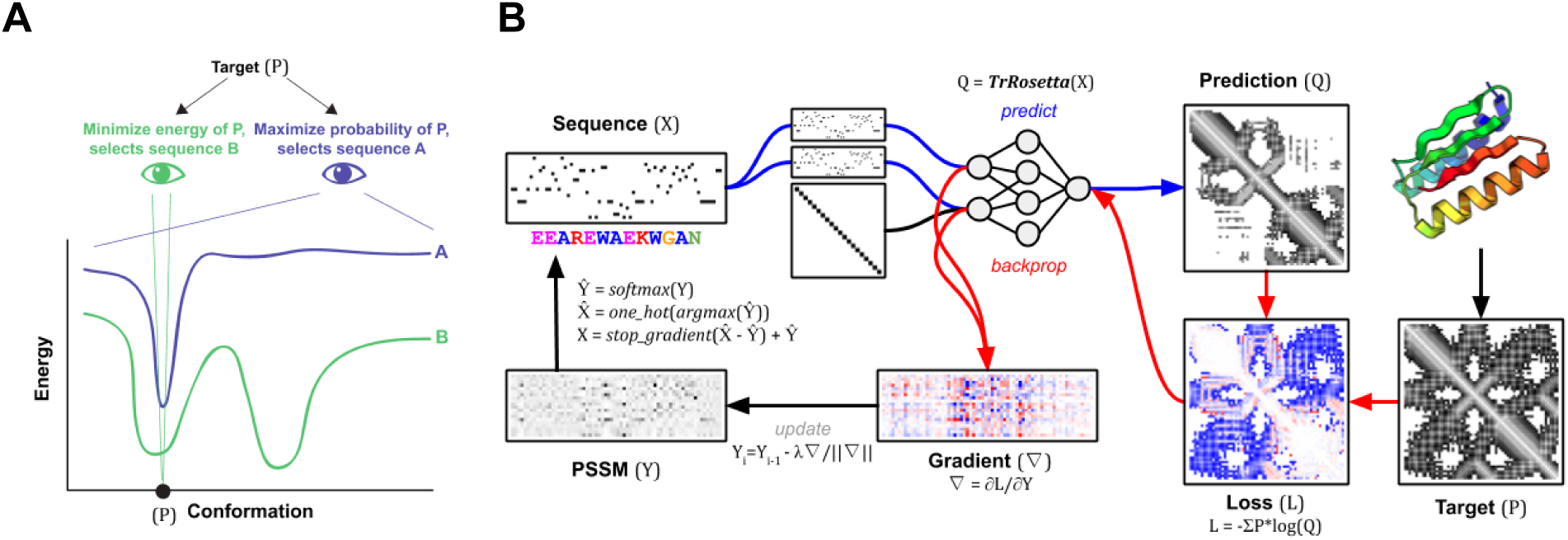
**(A)** The goal of fixed-backbone protein design is to find a sequence that best specifies the desired structure (P). Traditional energy-based methods have approached the problem heuristically; focusing solely on minimizing the energy of the target conformation in the hope that any stable alternative conformation is unlikely to arise by chance. Unfortunately, proteins designed by energy minimization, may adopt alternative conformations as suggested in the energy landscape of sequence B. An ideal method should instead find the sequence that maximizes the probability of the desired structure over all other states. Such a method would select sequence A. (**B)** Overview of trRosetta fixed-backbone sequence design method. Starting with a random profile (PSSM), the maximum value at each position is taken to generate a one-hot encoded sequence, which is fed into the trRosetta model. The output is the predicted distribution of distances, angles and dihedrals for every pair of residues (note, here we only show distances). The loss is defined as the difference between the target and prediction, and the gradient is computed to minimize this loss. The gradient, after normalization, is applied to the PSSM and the process is repeated for 100 steps or convergence.

## Results and Discussions

### Sequence design by gradient backpropagation

We set out to adapt trRosetta for the classic “fixed backbone” sequence design problem by developing a suitable loss function assessing the probability of the desired structure for a given sequence, and an efficient optimization method for finding sequences which maximize this probability. For the loss function, we simply sum the logarithms of the probabilities of the observed inter-residue distances and orientations – a proxy to the log-likelihood of the sequence given the structure. For optimization, we experimented with a simple iterative procedure in which a sequence is: *i*) randomly generated and input to the network *ii*) the gradient of the loss is backpropagated to the input sequence (treated as a *N x* 20 PSSM), *iii*) new amino acid sequence distributions are obtained at each position through a step down the gradient, *iv*) a single new amino acid sequence is obtained by selecting the highest probability amino acid at each position, *v*) this sequence is fed back into the network for the next iteration/update (Fig. 1B). We found that this procedure converged for a variety of ∼100-residue protein structures after ∼25 iterations, requiring only a few minutes of GPU time. We benchmarked this optimization method by redesigning 2000 proteins previously generated using MCMC optimization with trRosetta (14). For all cases, the gradient descent approach can find a similar or better scoring solution within a 100 iterations (Fig. S2; for comparison to a related backpropagation method (15), see Methods and Fig. S3).

In many design applications, it is desirable to generate not just one sequence which folds to a given structure, but an ensemble of sequences. In Rosetta, this is typically done through independent Monte Carlo sequence optimization calculations, which are CPU-time intensive. We reasoned that our trRosetta based sequence design approach could be extended to generate not just one sequence, but ensembles of thousands of sequences in one pass; by taking as the variables being optimized the identities of the amino acids of 10,000 aligned sequences. This is straightforward since trRosetta’s network already takes aligned sequences as inputs. As shown in Fig. S1, such “sequence alignment” design generates multiple sequence alignments with residue-residue covariation and other hallmarks of naturally occurring alignments. This information could be very useful in guiding the construction of smart sequence libraries for directed evolution of enzyme activities and other properties where the number of sequences in the naturally occurring family is too small to adequately estimate these features.

### trRosetta captures general properties of the folding energy landscape

Because trRosetta generates probability distributions over all possible structures, we reasoned that it might be able to detect overall properties of folding energy landscapes better than conventional methods such as Rosetta, which only “see” the target structure (Fig. 1A). To test this, we collected a set of 4,204 monomeric proteins designed by Foldit players, who compete to optimize the Rosetta energy of design structures without the aid of energy landscape calculations (16). We used *ab initio* folding calculations to probe their energy landscapes – generating tens of thousands of conformational samples (decoys) for each designed sequence – by examining the relationship between the energy of each decoy (as computed by Rosetta) and its structural deviation from the designed state. Some energy landscapes are characterized by sharply funneled profiles leading into the designed target structure, while others are relatively flat, with little or no energy gap between the designed structure and very different structures. These differences were quantified using an estimate of the Boltzmann probability of the target structure (*Pnear*: see Eq. 6 in Methods).

We found that trRosetta was a better predictor of designs with high *Pnear* scores than the Rosetta energy function (Fig. 2A). For energy landscapes with low *Pnear* values, the likelihood of the designed structure given the sequence was lower on average than for designs with high *Pnear* values. Hence, as hypothesized above, the trRosetta probability distributions assign higher probability to the reference (target) structure when the energy landscape is more funneled. Being able to assess energy landscape properties from a calculation on a single structure is quite remarkable, and should have considerable practical utility; energy landscapes characterization using Rosetta *ab initio* folding simulations are extremely computationally intensive, requiring thousands of CPU-hours to generate sufficient decoy structures to map out the landscape, while the trRosetta calculations takes seconds on standard GPUs. As energy landscape evaluation is the final step in *de novo* protein design before experimental characterization – to determine the extent to which the sequence encodes the structure – this much more rapid approach to evaluate landscapes should considerably streamline the *de novo* design process.

**Fig. 2.**
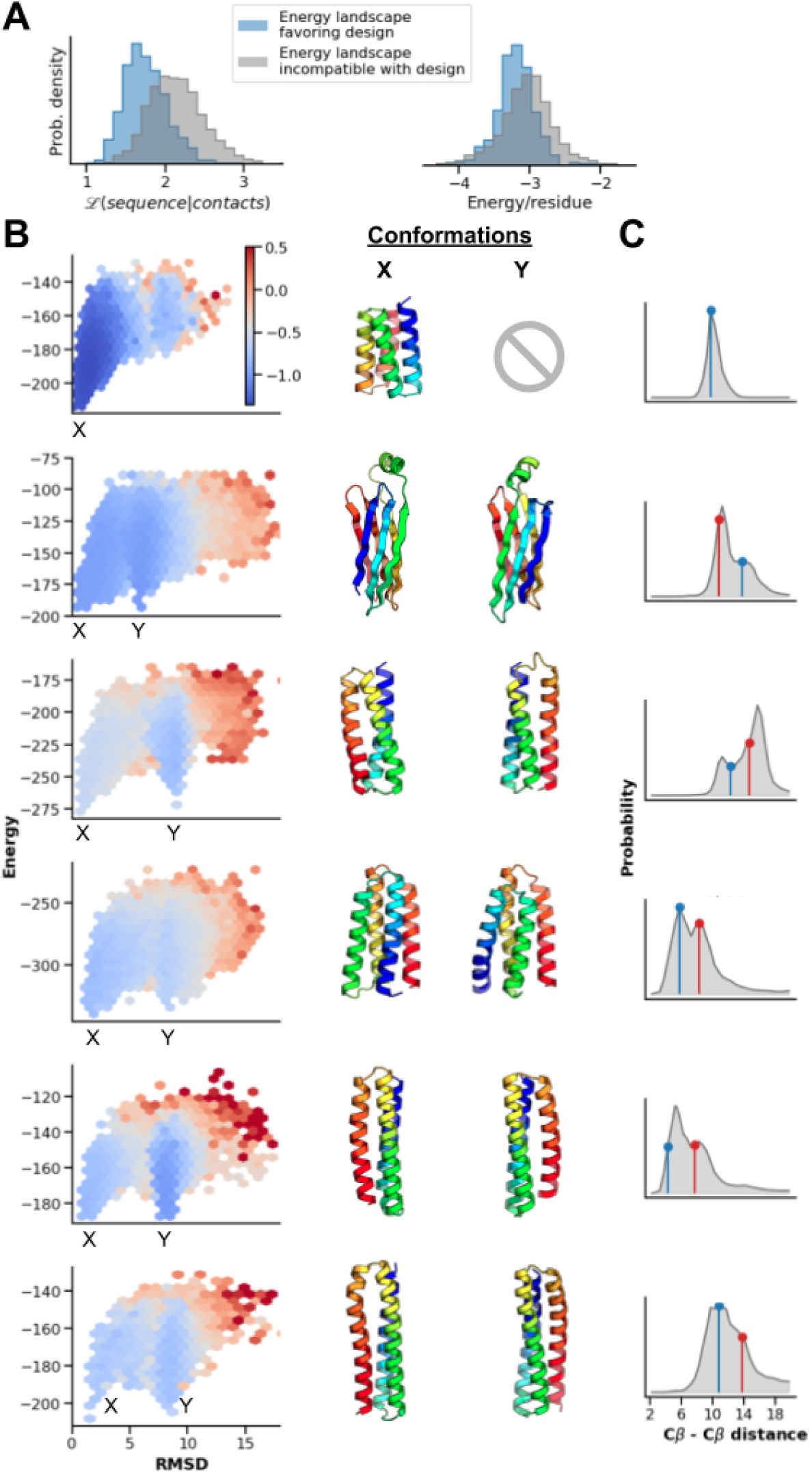
TrRosetta predicts properties of the folding energy landscape. (**A**) Unlike Rosetta-based energy calculations, which only see the target conformation, trRosetta better predicts which designs will have high Boltzmann-probabilities (*Pnear* > 0.8, *AUC*_*trRosetta*_ = 0.81 vs. *AUC*_*Rosetta*_ = 0.65, *N* = 4,204). (**B**) Relative to an energy landscape with just a single energetically favorable, and thus highly probable, conformation (first row), trRosetta correctly predicts dilution of probability mass for 5 designs with multiple low-energy conformations (rows 2-6). Structural decoys were binned on a hexagonal grid, and the mean trRosetta score (ℒ (*sequence*|*structure*), corrected for background, see Methods) represented by the color gradient. States X and Y are the lowest energy representatives. (**C**) Examples of probability distributions (*Cβ-Cβ* distance prediction) for specific *i,j* pairs. The actual distances in the X, Y conformations are shown by vertical lines (blue and red respectively).

Next we investigated how trRosetta distributes probability density across the energy landscape when alternative low-energy states are present. To assess this we computed the probability of each of the tens of thousands of decoy structures spread throughout the energy landscape with trRosetta, and compared the distribution of probability to those for energy landscapes strongly funneled into the designed structure. When no alternative energy minima were present, probability density was concentrated near the designed target structure, and decreased monotonically with increasing distance from the target structure in both the structural and energy dimensions (Fig. 2B, first row). In contrast, for designs with alternative minima, the total probability density of the designed target structure was lower, with substantial probability “leaking” into the alternative minima (Fig. 2B, rows 2-6). Thus, the trRosetta calculation of structure probability density from sequence recapitulates not only the Boltzmann probability of the design target structure, but also – in cases with alternative minima – the spreading out of this probability density into specific alternative states.

We investigated how the trRosetta predicted probability distributions encode the existence of multiple structures for one sequence. Examination of the probability distributions for specific residue-residue distances for energy landscapes with multiple minima revealed that for some *i,j* pairs the distributions were bimodal (Fig. 2C and Fig. S4), with one peak corresponding to the design target state, and the second corresponding to the alternative minimum. This suggests that trRosetta calculations from sequence may be recognising the presence of alternative states *explicitly*. In other cases, the maximum predicted distance probability was in between the two low energy states; distance features may get averaged and broadened in cases where the model is less certain about the structure (see Fig. S4). These observations have implications for multistate design; probabilistic models such as trRosetta may enable the simultaneous optimization of two or more structures – a challenging task for energy-based design methodologies.

### trRosetta identifies more stable *de novo* designs

As a first step towards using trRosetta for *de novo* protein design, we took advantage of a high throughput protease resistance based protein stability assay that enables experimental quantification of the stability of tens of thousands of designed proteins in parallel. In addition to 16,174 already characterized mini-proteins within the topologies HHH, HEEH, EEHEE, and EHEE (17), we designed a new set of 13,985 sequences that fold into four different β-sheet topologies using Rosetta. Genes encoding these designs were encoded in large oligonucleotide arrays, transformed into yeast cells, and the encoded designs displayed on the surface of the cells. Treatment with trypsin and chymotrypsin at different concentrations followed by fluorescence activated cell sorting and deep sequencing was used to quantify the stability of each design. We then determined the probability of each designed structure given the designed sequence according to trRosetta, and investigated the extent to which this distinguished stable designs from unstable designs. As shown in Fig 3A, the trRosetta probability calculations indeed distinguished stable from unstable designs, with designs with high probability structures given their sequence being more stable on average than designs with low probability. The trRosetta probability was a better predictor of stability than the Rosetta energy (Fig 3A, right, and Fig. S5).

**Fig. 3.**
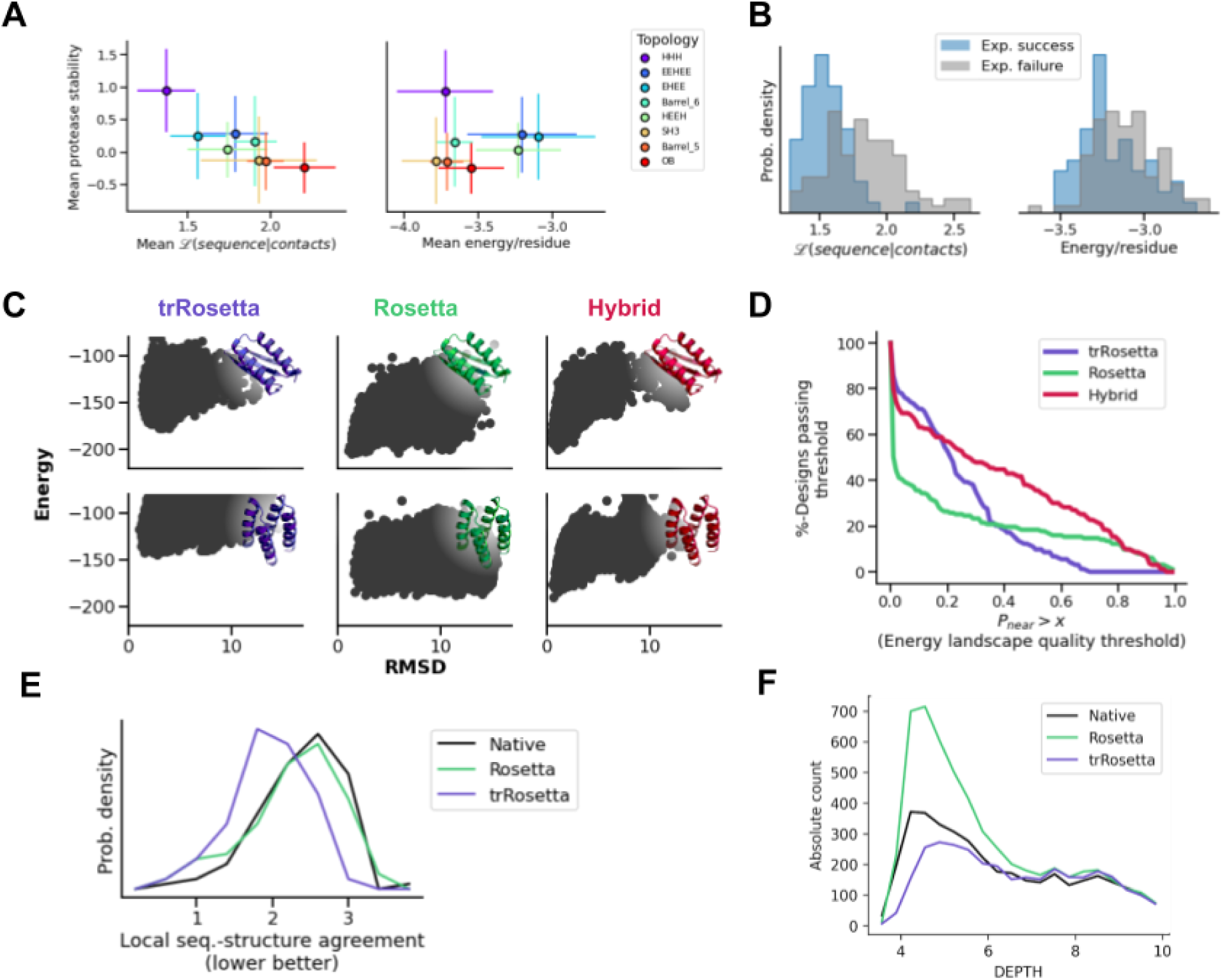
(**A-B**) TrRosetta predicts scaffold designability and experimental success. (**A**) Across different topologies, trRosetta predictions are better correlated with experimental protease stability – a measure of folding success – than Rosetta energy (*R*^*2*^ _*trRosetta*_ = 0.79 vs. *R*^*2*^ _*Rosetta*_ = 0.00). Datapoints are the topology-specific mean values, and error bars represent standard deviations (8 topologies, *N*_*tot*_ = 30,159 designs). (**B**) TrRosetta is significantly better at discriminating experimental success (expression, non-aggregation, and having correct secondary structure content) than Rosetta energy (*AUC*^*trRosetta*^ = 0.81 vs. *AUC*_*Rosetta*_ = 0.64). Data from 145 Foldit-generated designs (29). (**C-D**) Designing with a hybrid trRosetta/Rosetta protocol disfavors off-target states. (**C**) Examples of energy landscapes for two Foldit-generated backbones, each designed with trRosetta, Rosetta, and the trRosetta/Rosetta hybrid protocol. (**D**) The hybrid protocol improves the quality of the resulting energy landscapes, as determined by the *Pnear* quantity. TrRosetta on its own also improves funnels, but only superficially (better performance than Rosetta in the lower *Pnear* regime). It does however not generate deep minima in the vicinity of the designed state (poorer performance than Rosetta in the high *Pnear* regime). (**E)** The local sequence-structure relationship is idealized for trRosetta designs compared to both native proteins and Rosetta designs. The native structures that were used for re-design were at most 30% sequence identical to any protein in the trRosetta training dataset. Local sequence-structure agreements were measured as the average RMSD between the designed structure and 9-residue fragments from the PDB that were selected based on the sequence of the design. **(F)** For the same set of native backbones, trRosetta re-designs have a more native-like distribution of hydrophobic residues (F,I,L,V,M,W,Y) on the protein surface than Rosetta re-designs. The degree of burial was assessed with DEPTH (30). For breakdown by amino acid, see Fig. S12.

In laboratory experiments, designed proteins can fail for a variety of reasons, even after passing stringent computational selection criteria. Typical problems include lack of soluble expression, aggregation, and folding into unintended structures. Reasoning that many of these problems could be associated with poor energy landscapes, we evaluated the ability of trRosetta to predict the experimental success of 145 Foldit-player proteins that had been selected for experimental testing based on *ab initio* folding (16). Compared to Rosetta energy, trRosetta better predicts whether the protein can be expressed, purified and is folded (Fig. 3B). trRosetta also had considerable predictive power for expression alone, aggregation, and monomericity (Fig. S6). While more extensive characterization on a wider variety of designed proteins will be important to establish generality, given that experimental characterization is a bottleneck in protein design, trRosetta could become a useful tool for pre-filtering designs.

### Sequence design using trRosetta disfavors off-target states

The landscape awareness encoded in trRosetta’s probabilistic description of sequence-structure relationships suggests that it could be used to design sequences that maximize the probability of the desired state explicitly, by avoiding the presence of alternative states. To investigate this possibility, we re-designed a diverse set of backbones with the backpropagation method described above; we grouped the 4,204 Foldit designs into 200 structural clusters spanning a large range of topologies (Fig. S7), and picked one structure from each cluster. The same backbones were also re-designed with Rosetta (see Supplementary Methods), providing a direct comparison of the consequences of minimizing energy *versus* maximising probability. Following design, large scale *ab initio* folding calculations were performed to map out the energy landscape of each sequence, and the occupancy of the target structure computed as described above.

The energy landscapes of sequences designed with trRosetta had on average higher *Pnear* values than those of sequences designed with Rosetta (Fig. 3C and Fig. S8), consistent with the fuller view of the energy landscape. However, in some cases, the predicted energy landscapes had less pronounced funnels leading into the target structure, and the energy gap between the target structure and the lowest lying alternative states was smaller. This likely reflects the relatively low-resolution representation of the structure used by trRosetta: accurate energy calculations require sub-angstrom resolution, which the angle/distance bin-sizes of trRosetta do not capture. Another likely consequence of this low-resolution representation, is trRosetta’s limited understanding of the thermodynamic effect of single-point mutations (Fig. S9). Thus, while trRosetta is better at capturing the global features of energy landscapes, the Rosetta full-atom description is more capable at creating deep energy minima at the target structure. These results suggest that trRosetta is more adept at disfavoring alternative minima, and Rosetta, at creating a deep minimum at the target structure: in a simple energy landscape picture, trRosetta may be viewed as smoothing the landscape away from the minimum, and Rosetta, as making the minimum as deep as possible.

As the two approaches have different strengths, we hypothesized that a combined method might prove more effective than either alone. We evaluated the performance of a hybrid trRosetta-Rosetta design protocol by redesigning the same set of backbones, followed by *ab initio* folding calculations. For each target structure, trRosetta was used to generate a PSSM (by designing one hundred unique sequences), which were then used to constrain the set of amino acids allowed at each position in Rosetta sequence design calculations (see Supplementary Methods). This approach proved superior to either method on its own, leading to more funneled energy landscapes (Fig. 3D, red line), attesting to the strength of combining design methodologies working at different levels of resolution. This hybrid method also outperformed two state-of-the-art design protocols (18, 19) that attempt to compensate for the lack of energy landscape awareness of Rosetta design calculations (Fig. S10). We anticipate that this hybrid method will be broadly useful, as it combines a moderate resolution representation of the overall energy landscape with a very detailed atomistic representation around the designed target structure.

What properties of folding free energy landscapes are captured by trRosetta? Physical interpretation of representations within neural networks with millions of parameters is not straightforward, but we can identify at least three distinct contributions. First, the trRosetta network likely identifies alternative global and supersecondary structure packing arrangements (Fig. 2B), which are reflected in the bimodal distance distributions (Fig. 2C). Second, designs made with trRosetta have more ideal local sequence-structure relationship compared to Rosetta-based designs and native proteins (Fig. 3E). Third, trRosetta designs fewer hydrophobic residues on the surface of proteins than Rosetta (when not explicitly restricted) (Fig. 3F); surface hydrophobic residues do not appreciably change the energy of the designed minimum, but can stabilize alternative structures in which these residues are buried.

## Conclusions

Our results demonstrate that sequence design using trRosetta has the remarkable ability to capture properties of the entire energy landscape and consider alternative states that can reduce the occupancy of the desired target structure. Such implicit considerations of the full landscape are almost impossible to achieve with atomistic models without employing extremely CPU-intensive calculations, like the large scale Rosetta *ab initio* structure predictions employed here, or molecular dynamics simulations on very long time-scales. On the other hand, because of the lower resolution of the trRosetta method, it is less accurate in the immediate vicinity of the folded structure. Our integration of trRosetta design into Rosetta all-atom calculations appears to combine the strong features of both approaches, and we expect should be broadly useful. More generally, this work demonstrates how deep-learning methods can complement detailed physically based models by capturing higher-level properties normally only accessible through large scale simulations.

## Methods

### Approach

The fixed backbone protein design problem is to find an amino acid sequence compatible with a target structure. Probabilistically, one seeks for a sequence which maximizes the conditional probability *P(sequence*|*structure)*. Using Bayes theorem the sought-for probability can be equivalently expressed as

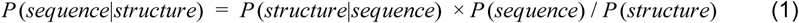

In trRosetta, the structure is represented as a tensor of 6D transformations between every pair of residues within 20 Å (*Cβ-Cβ* distance). Since the protein structure is given and does not change in the course of design (*P(structure) = const*), the design problem is equivalent to maximizing the *P(structure*|*sequence) × P(sequence)* product on the right-hand side of Eq. 1. To approximate the *P(structure*|*sequence)* term, we use a pre-trained trRosetta structure prediction network (13) which predicts residue-residue distances and orientations from an input sequence or a multiple sequence alignment. In the reverse problem of fixed backbone protein design, the structure is given, while the sequence is variable, so the quantity to be optimized (*ℒ(sequence*|*structure)*) can be viewed as the likelihood of a sample sequence given the protein backbone.

For every residue pair (*i,j*), trRosetta generates probability distributions over *Cβ-Cβ* distances *p(d*_*ij*_) and orientations described by three dihedrals (*p(*ω_*ij*_*), p(θ*_*ij*_), and p(θ_ji_)) and two planar angles (*p(φ*_*ij*_) and *p(φ*_*ji*_), see (13) for details). The likelihood is then computed from the network predictions as the average over all residue pairs and all coordinates *y* ∈ {*d*, ω, θ, φ} at coordinate values *y*^0^ derived from the input structure:

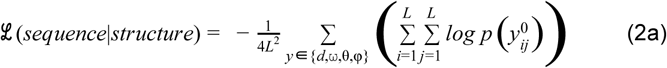

We found it often helpful to limit the likelihood calculations to residue pairs actually in contact (*Cβ-Cβ <* 10 Å) in the target structure, yielding ℒ (*sequence*|*contacts*).

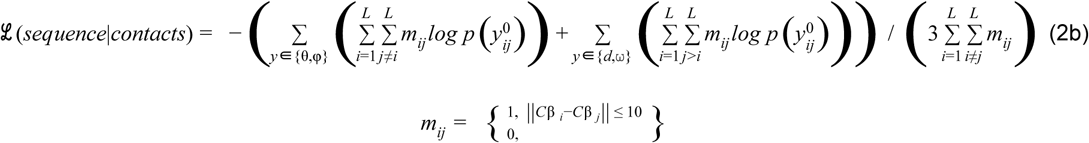

Under certain circumstances, it can be useful to ‘normalize’ predictions to allow side-by-side comparison to be made. In this case, the probability (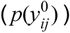)) in Eq. 2a can be replaced by:

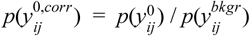

where 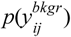 is the background probability, which was obtained by passing sequence-agnostic input features represented by random gaussian noise to a separate network with similar architecture to trRosetta. These background inter-residue probabilities can be interpreted as ‘average distributions’ across the entire PDB. This correction was applied to the energy landscape plots shown in Fig. 2B.

For *P(sequence)* we use the amino acid composition biasing term described in (20), so the total loss takes the form:

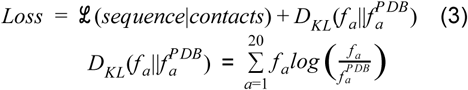

and Eq. 3 is subject to minimization with respect to the input amino acid sequence. In cases where the average amino acid composition of the PDB is not desired, an amino acid specific bias loss can be computed for specific tasks such as native sequence recapitulation (see supplementary methods).

### Optimizing the loss with gradient descent

The architecture of the trRosetta network allows for computing gradients of the network outputs with respect to its inputs by backpropagation (13). This means that Eq. 3 can be optimized by simple gradient descent. Since the input to the network is a discrete variable (an amino acid sequence) one also needs a proper way of tweaking the sequence in response to the calculated gradient. To this end, we introduce a continuous random variable *Y*_*Lx20*_ initialized normally, then normalize each row *Y*_*i*_ separately with the *softmax* function *softmax(Y*_*i*_), and finally choose the entry with the highest value so that

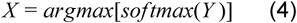

where both *argmax* and *softmax* are only applied to the second dimension of the argument. Modifying network inputs according to Eq. 4 makes it possible to directly optimize the loss in Eq. 3 with respect to the auxiliary variable *Y* (21):

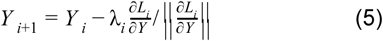

To have more control over the minimization, in Eq. 5 we normalize gradients (22)) and gradually decrease the step size λ_*i*_ according to the non-linear schedule λ_*i*_ = ((1 − *i*) / *N*) ^2^ where *N* is the number of minimization steps.

A similar approach using backpropagation through the trRosetta network was recently described (15); the two key differences include the normalization scheme and the loss function. Instead of normalizing and sampling from the PSSM, we normalize the gradients and take the argmax of the PSSM. Instead of minimizing the KL divergence, we minimize the categorical cross entropy. For the same task and loss, we find our optimization scheme requires significantly fewer steps (50 vs. 1000) to achieve the same loss (Fig S3).

### *Ab initio* folding calculations

Energy landscapes of designs were mapped out using Rosetta *de novo* structure prediction (*ab initio* folding) (23). In brief, short structural fragments from the PDB are collected using a bioinformatic pipeline taking the designed sequence as input. Next, starting from an extended conformation, the designed sequence is ‘folded’ by insertion of these fragments (substitution of the backbone torsion angles by that of the fragment) using a Monte Carlo simulated annealing protocol minimizing the energy. Thousands of such trajectories are run on Rosetta@home, generating different decoy conformations for the queried sequence. The energy landscape is represented by plotting the structural deviation between the decoys and the designed structure against their energies.

The quality of the energy landscape was quantified using the *Pnear* quantity (24), which approximates the Boltzmann-weighted probability of finding decoys near the designed state (fuzzy cutoff):

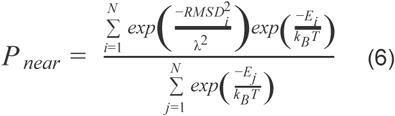

where *N* is the number of decoys, *E* is the energy of the decoy, and *RMSD* represents its structural deviation from the target state. The stringency for nativne-ness is controlled by *λ* (set to 3), and the temperature factor (*k*_*B*_*T* = 0.62) controls the sensitivity of the score to low-energy alternative states. The *Pnear* value ranges from 0 (energy landscape incompatible with the designed state) to 1 (energy landscape favoring the design).

### Prediction of protease stability

We used trRosetta to predict stability on a dataset composed of the designs from Rocklin *et al*. (25) (*N* = 16,174, four different topologies: HHH, HEEH, EHEE, EEHEE) and 13,985 small β-barrels designs (four different topologies: OB, SH3, Barrel_5, Barrel_6). Backbones were constructed using blueprints (26), followed by sequence design with Rosetta (FastDesign (24, 27) in conjunction with LayerDesign (18), and using the Rosetta all-atom energy function (28) with beta_nov16 weights). We performed the analysis across the entire dataset, as well as within each topological group, and compared the prediction to Rosetta energy (all designs were re-scored with beta16_nostab weights to enable comparisons).

### Prediction of experimental success

The experimental characterization of 145 Foldit player-designed proteins was reported previously (16). This dataset was used to assess trRosetta’s ability to predict experimental outcomes. Expression and solubility were assessed by SDS-PAGE; oligomeric state by size exclusion chromatography; and secondary structure content by circular dichroism.

## Acknowledgments

C.N. is supported by Novo Nordisk Foundation grant NNF17OC0030446. B.I.M.W. is an EMBO long-term fellow (ALTF 139-2018). S.O. and S.L. are supported by NIH grant DP5OD026389. D.K. and D.B. are supported by the Howard Hughes Medical Institute. B.K. and D.B. are supported by the Audacious Project at the Institute for Protein Design. I.A. is supported by the National Institute of Allergy and Infectious Diseases (NIAID, Federal Contract HHSN272201700059C). The authors are grateful for funds provided by Eric and Wendy Schmidt, by recommendation of the Schmidt Futures program. The source code for the study is available at https://github.com/gjoni/trDesign.

## Supplementary Figures

**Fig. S1.**
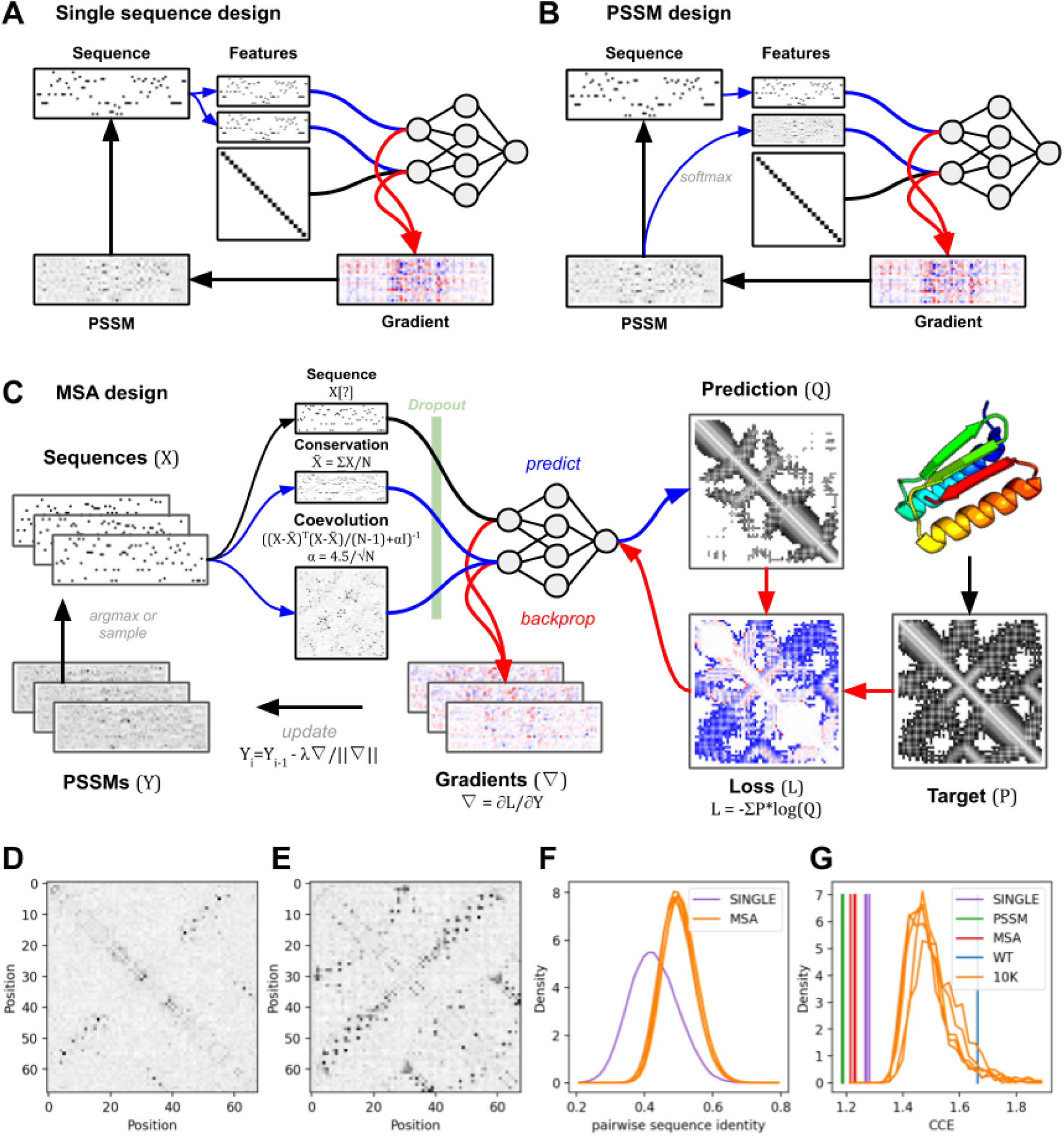
The inputs to trRosetta include the tiled one-hot encoded query sequence, conservation and coevolution matrix. For single sequence design (**A**) the conservation channel is substituted by sequence and coevolution channel with an identity matrix. For PSSM design (**B**), the conservation channel is the softmax of the PSSM. Finally for MSA design (**C**) conservation is the mean of all the sequences, and coevolution is the inverse covariance of the sequences. To prevent overfitting on the first sequence or any specific features, we only backpropagate through the conservation and convolution channels, randomly selecting a different sequence for the sequence channel, and included a dropout layer (80%) after tiling. (**D**) We often find that only a subset of contacts coevolve for independently generated sequences, while for MSA design, (**E**) all contacts have a covariance signal. (**F**) The diversity of designed sequences (pairwise sequence identity) and (**G**) cross-entropy scores for: Single sequence design, MSA design, PSSM design and [W]ild[T]ype sequence for a given backbone (PDB:6MRR). The orange distributions are the CCE (also termed ℒ (*sequence*|*structure*)) scores of the 10,000 MSA designed sequences when scored using the single-sequence predictor from (**A**).

**Fig. S2.**
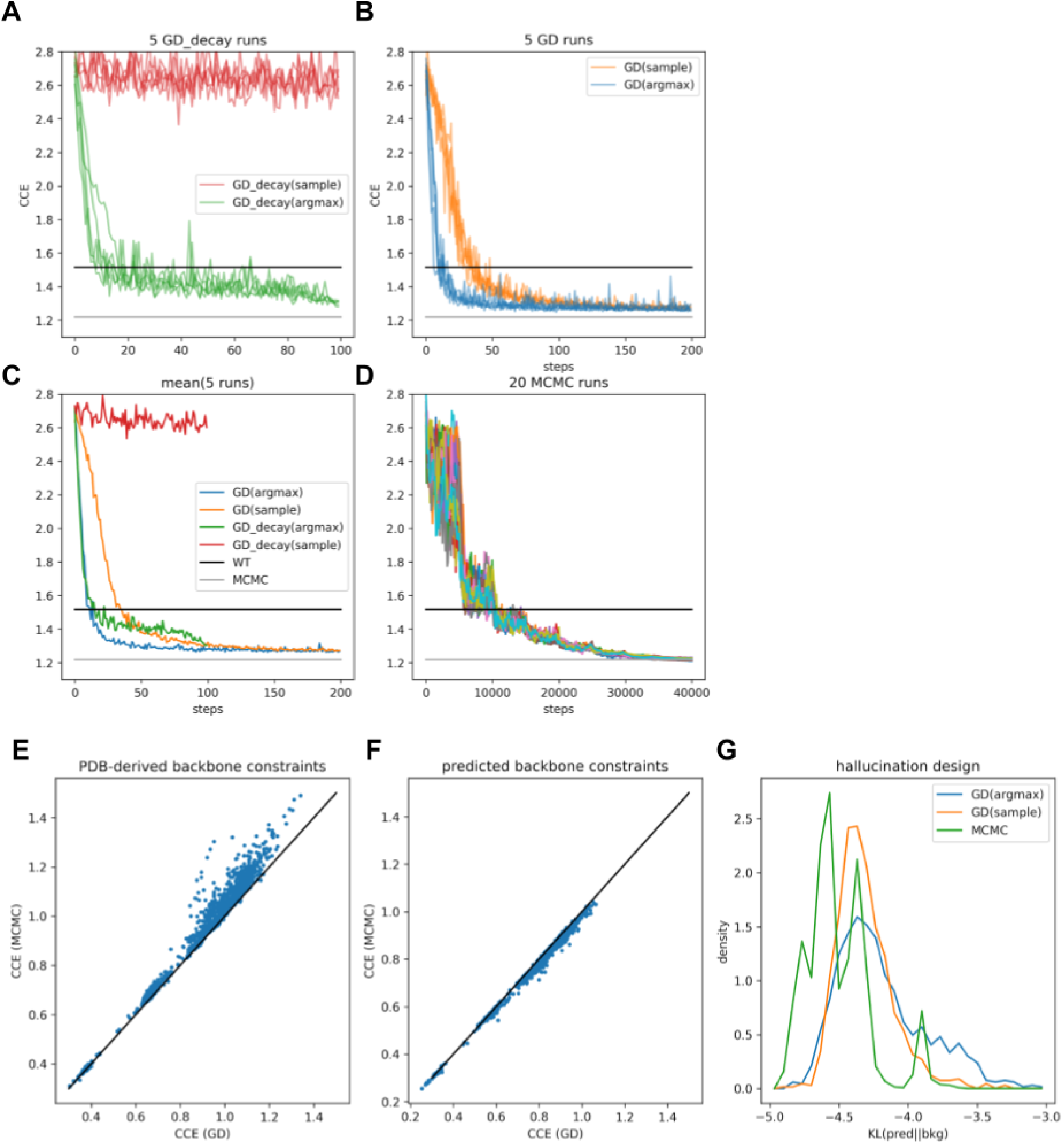
Comparing gradient descent (GD) based optimization to MCMC. (**A-D**) Optimization trajectories for Top7 (PDB:1QYS) fixed backbone design. (**A**) For decay-based minimization (as described in the main text), we find that taking the argmax (green-line) as opposed to sampling (red line) from the categorical distribution works significantly better, although if we disable the decay, and instead set a constant learning-rate of sqrt(length) (**B**), sampling eventually leads to a similar optimum. The black line indicates the CCE of the original sequence, and the gray line the average CCE at the end of the MCMC trajectories for the same backbone. (**C**) The mean of the four optimization methods. (**D**) 20 independent MCMC runs. (**E-G**) We compare GD to MCMC on a larger set of proteins designed with the MCMC protocol (Anishchenko *et al*. (14)). For (**E**) we design a new sequence to match constraints derived from the rosetta-relaxed backbone. For all cases, the GD method is able to find a sequence that is similar or lower (in terms of loss CCE) than what was hallucinated by trRosetta using the MCMC approach. (**F**) If we instead try to design a sequence to match the predicted backbone constraints of the hallucinated design, GD is able to find an alternative sequence to match these constraints with similar or slightly worse CCE. (**G**) For newly hallucinated designs, the MCMC protocol is able to sample sequences with lower KL divergences. Interestingly, GD with sampling enabled more frequently generates sequences that have lower KL divergences than with argmax.

**Fig. S3.**
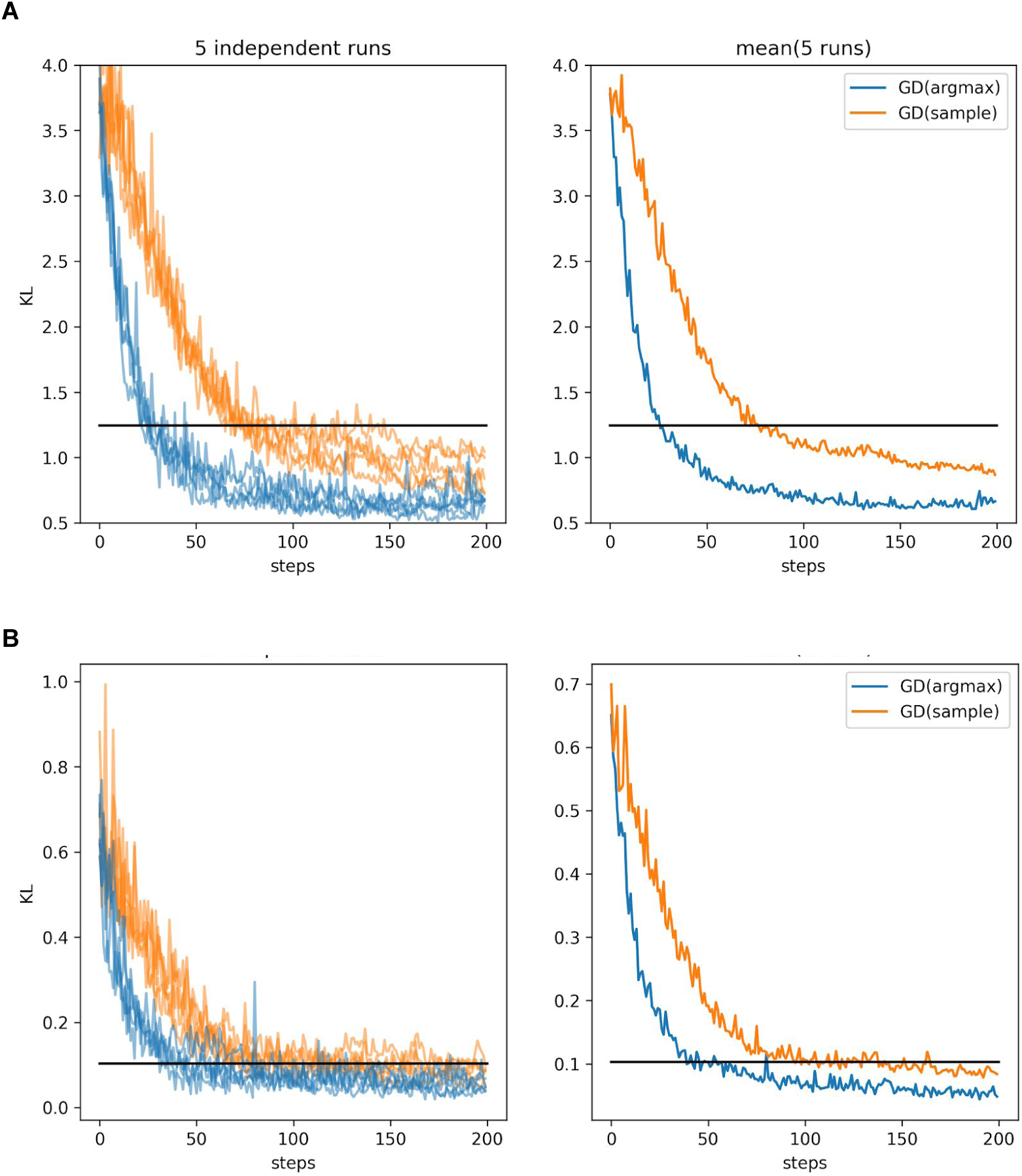
Testing our gradient descent (GD) based optimization on objective described in Linder *et al*. (31). (**A**) Optimization trajectories for KL(target||pred), where the target is the predicted constraints for Sensor Histidine Kinase protein. Within 50 steps, our protocol finds a sequence with a minimum lower than that reported after 1000 steps (black-line). (**B**) Here we rescore each sequence in the trajectory using a smooth KL (as was done by Linder *et al*.).

**Fig. S4.**
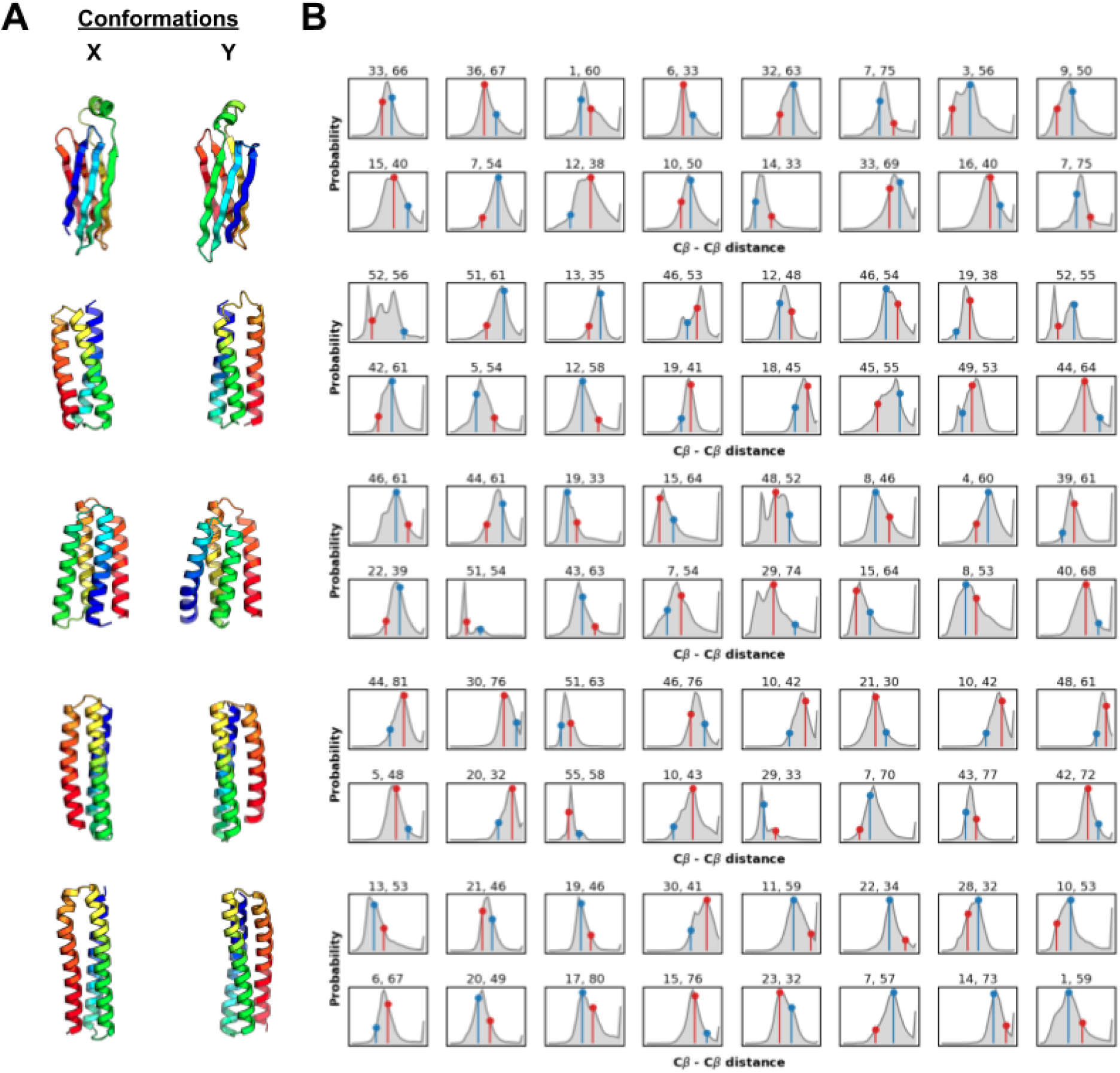
Examples of distograms from trRosetta for sequences adopting alternative conformations. (**A**) Structures of the Foldit-player designs that adopt alternative conformations as determined by *ab initio* folding (same as in Fig. 2B). The lowest energy states are shown (X; from the target funnel, Y; from the alternative state funnel). (**B**) Examples of outputs from trRosetta (*Cβ-Cβ* distance predictions) for specific *i,j* pairs (indicated above each plot). The actual distances from the target structure and the alternative state are show in blue and red respectively.

**Fig. S5.**
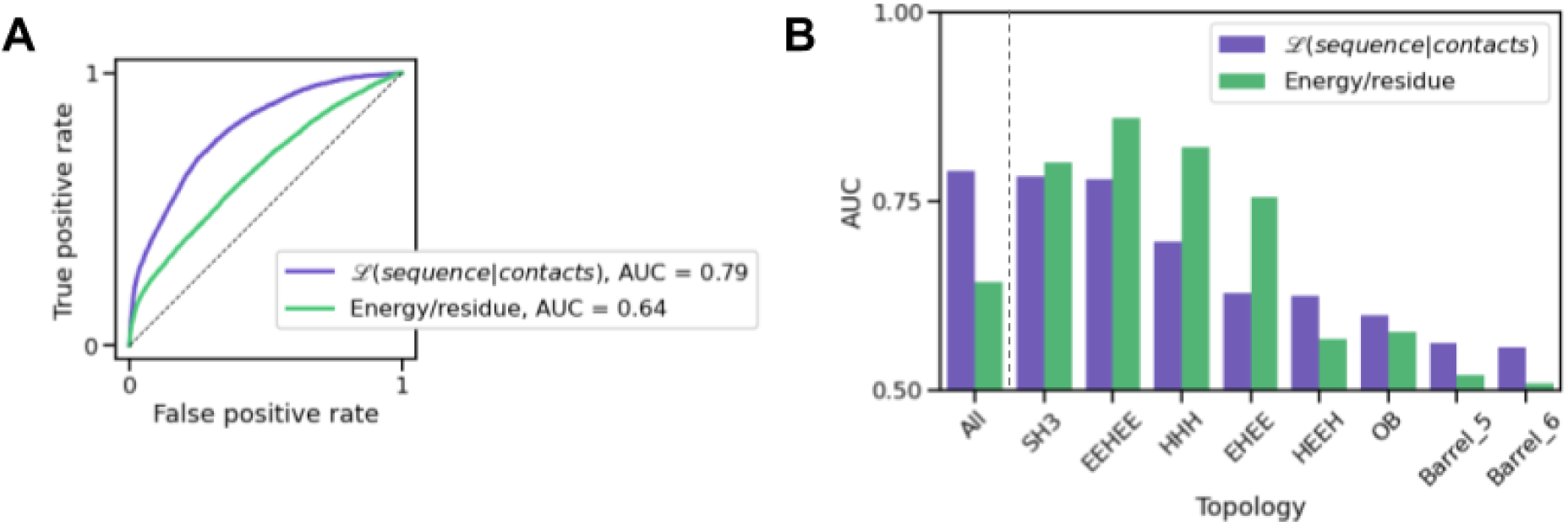
TrRosetta recognises protein stability across topologies, but Rosetta is often better at classifying intra-topology differences. TrRosetta (ℒ (*sequence*|*contacts*), purple) and Rosetta (energy/residue, green) scores were used to predict protease resistance (a proxy for folding stability, threshold = 0.5) for 30,159 designs spanning 8 topologies. (**A**) Receiver operating characteristic (ROC) curves showing the classification of protease stability data across all topologies. Area under the curve (AUC) values are indicated in the legend. (**B**) Comparison of AUC values for tRosetta and Rosetta classifications of stabilities across all designs (left of the dashed line, same as A), and *within* topologies (right of the dashed line).

**Fig S6.**
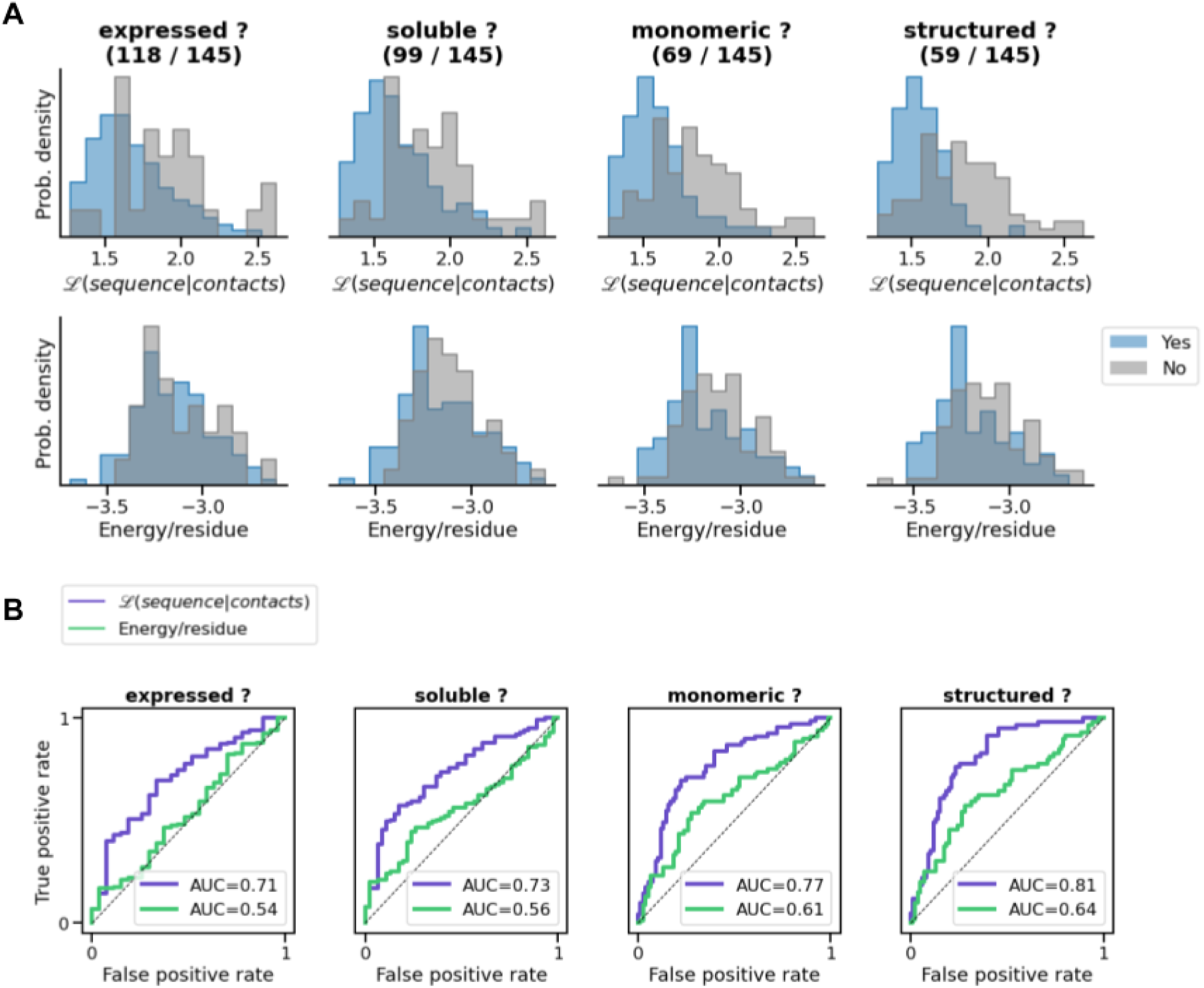
TrRosetta score is better at classifying experimental success than Rosetta. For 145 Foldit-player designs that were experimentally characterized (29), success was assessed at different stages: protein expression, solubility, oligomeric state, and presence of secondary structure after purification. (**A**) Histograms showing the distributions of success/failure at each experimental stage as assessed with trRosetta (top) and Rosetta (bottom). (**B**) The same data represented as ROC curves. AUC values for trRosetta (purple), and Rosetta (green) are shown on the plots.

**Fig S7.**
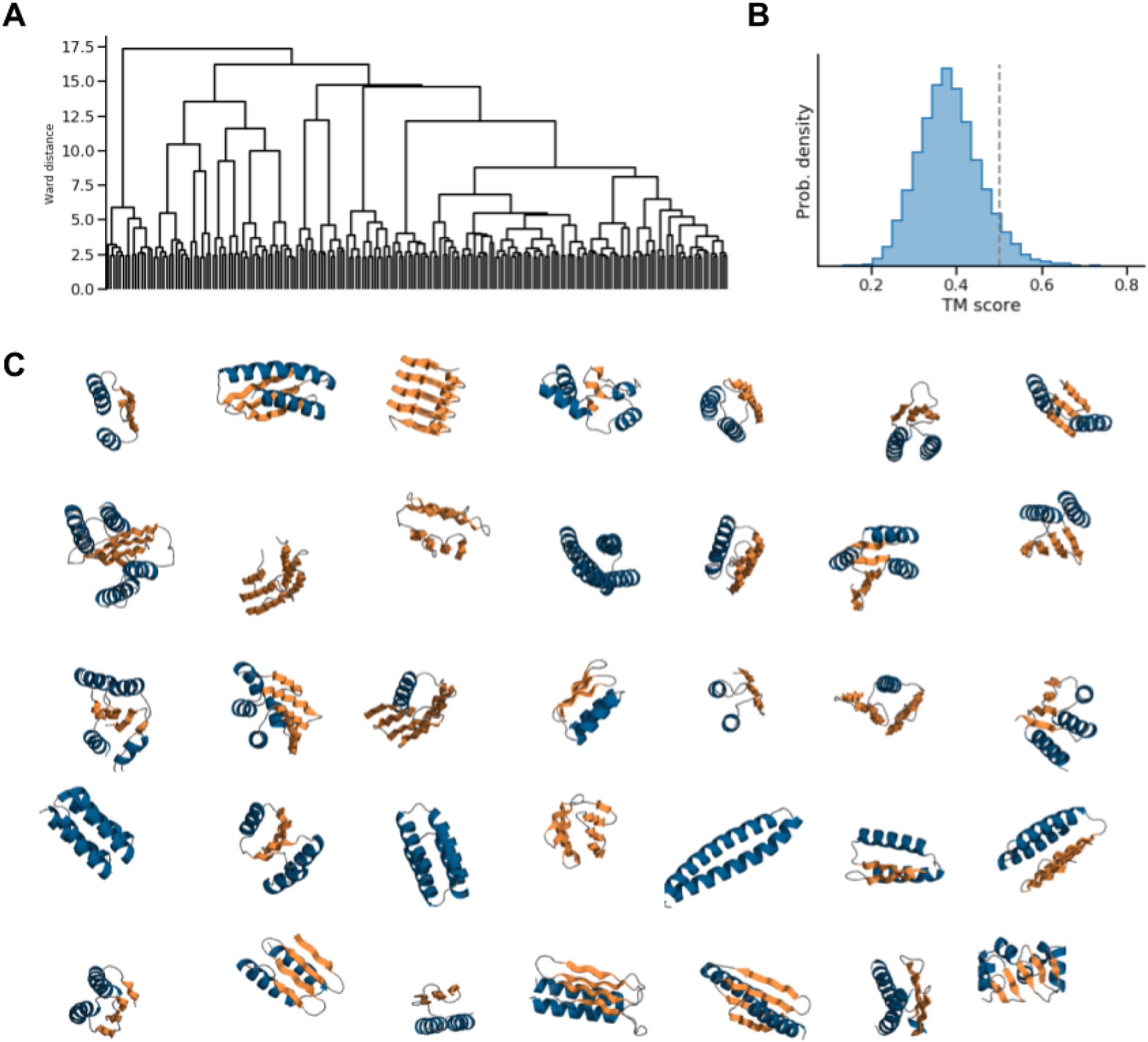
Structural clustering of Foldit players designs. (**A**) 5288 designs were clustered into 200 bins based on their pairwise TM-align scores (32). One design per bin was used for re-design. (**B**) Distribution of pairwise TM-align scores for the 200 selected designs (mean = 0.38) shows that they are structurally diverse. The threshold for homology (0.5) is indicated by a dashed line. (**C**) Examples of structures from the 200 selected designs, spanning all-α, all-β, and mixed α/β folds.

**Fig S8.**
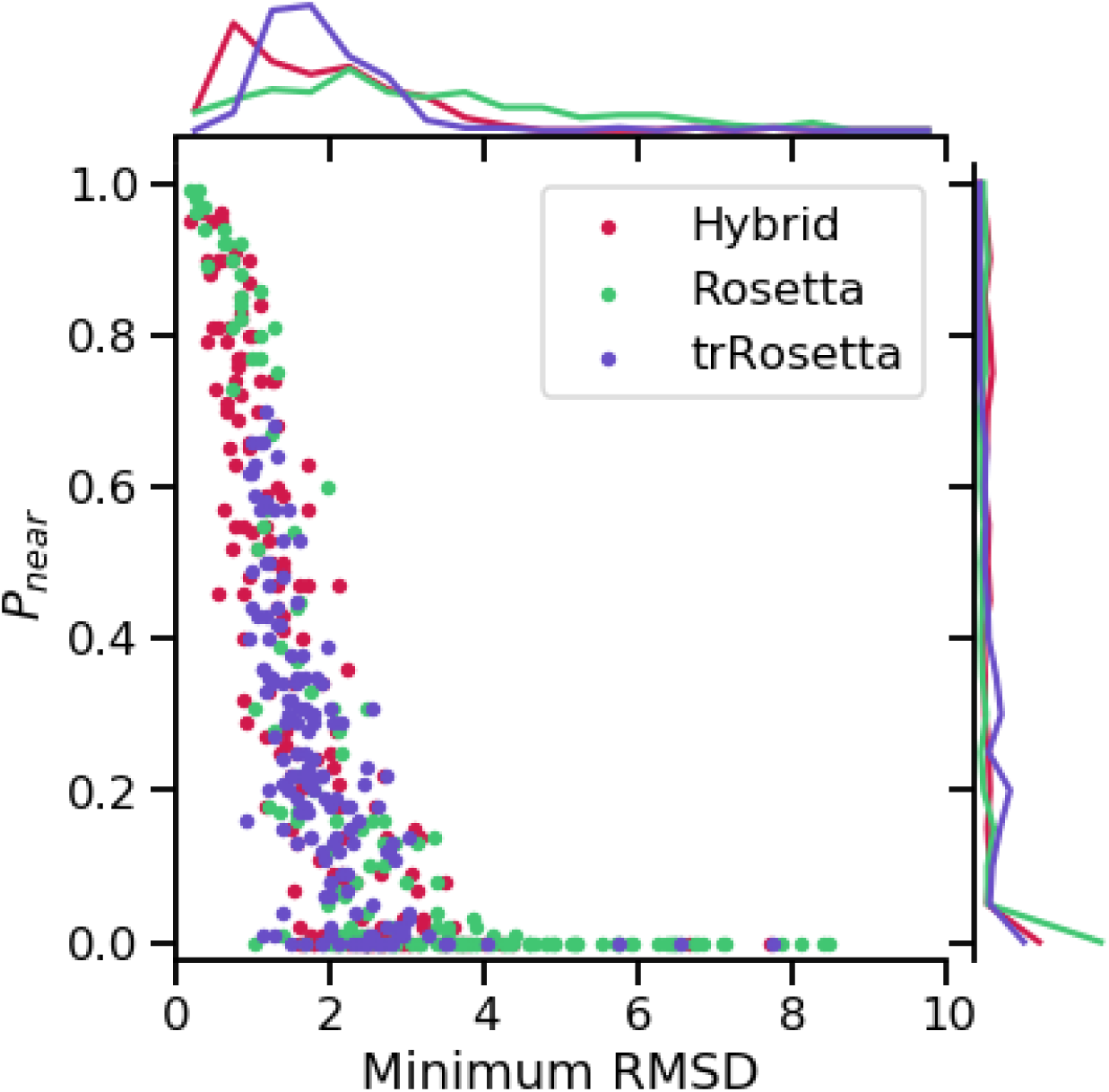
TrRosetta improves sampling of conformations near the target conformation. The energy landscape quality (*Pnear*) and RMSD of the lowest RMSD decoy were computed for the folding energy landscapes of 165 backbones designed with trRosetta, Rosetta or the Hybrid (trRosetta PSSM + Rosetta) method. In contrast to trRosetta-based methods (trRosetta and Hybrid), decoys from *ab initio* folding of Rosetta-designed sequences rarely sampled the target backbone conformations.

**Fig S9.**
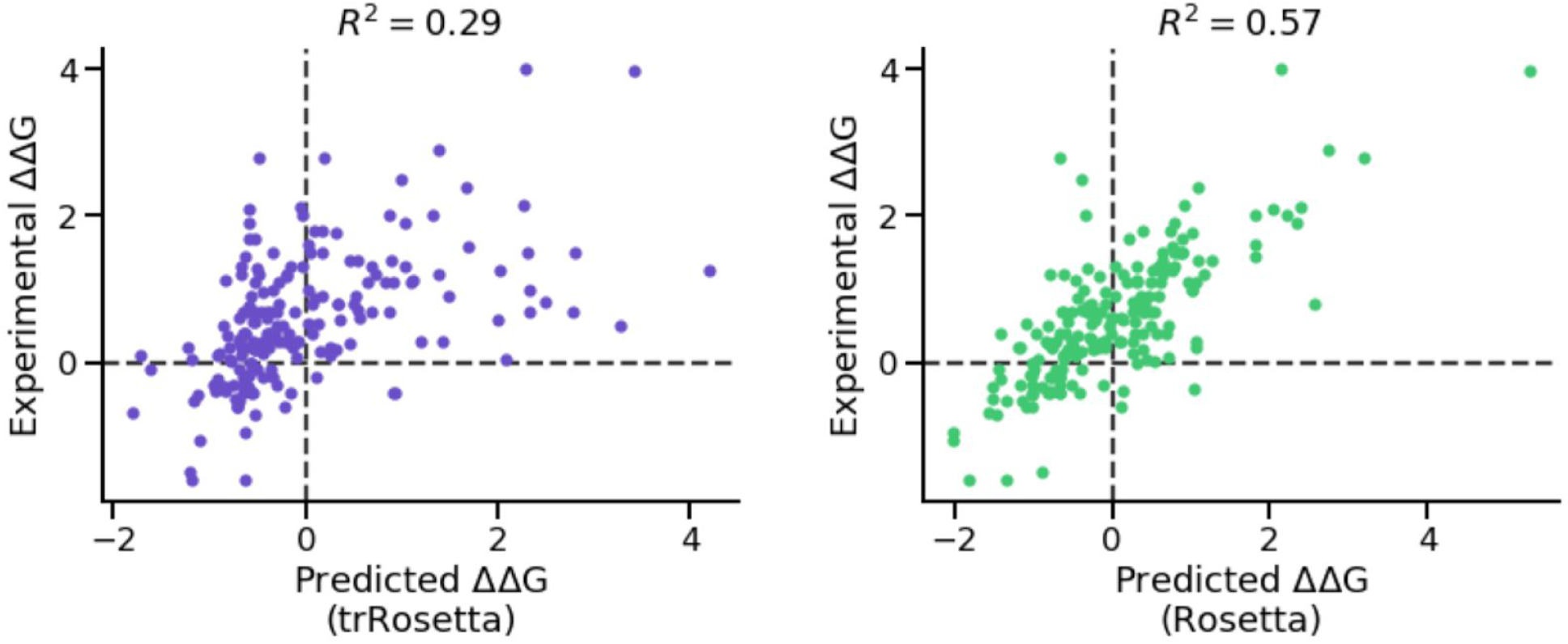
The Rosetta all-atom energy function (right) is better at predicting the thermodynamics of single point mutations than trRosetta (left). Based on a curated set of single point mutations from (33), we sub-selected the 15% best-scored sites with respect to ℒ (*wildtype sequence*|*mutation environment contacts*), where *mutation environment contacts* is defined as the subset of *i,j* pairs encompassing the contacts between the mutated position and the positions within 10 Å of it (*Cβ-Cβ* distances). The effect of mutations was computed as the difference ℒ (*wildtype sequence*|*mutation environment contacts*) -ℒ (*mutated sequence*|*mutation environment contacts*). Predicted Roseta ΔΔGs were taken from (34).

**Fig S10.**
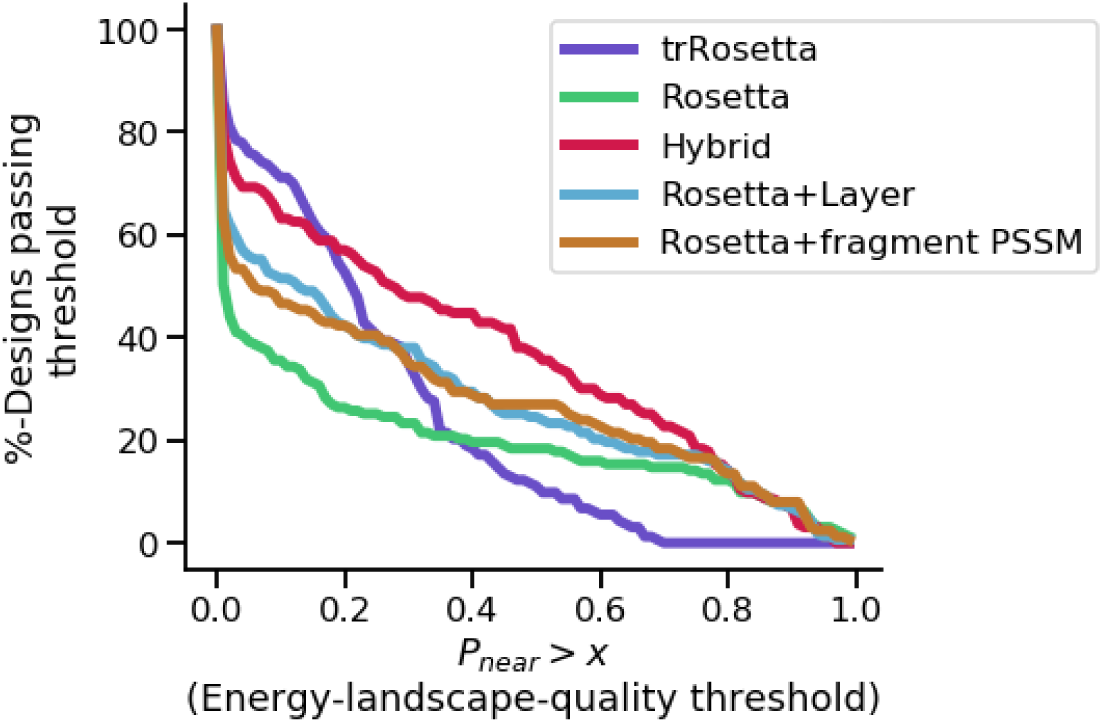
Performance curves for various sequence design methods. Compared to Rosetta design (all amino acids possible at all positions), restricting amino acid choices by burial (LayerDesign) or fragment-based PSSMs improves the mean energy funnel quality, but not beyond the quality of the Hybrid method, which applies trRosetta-based PSSMs to restrict and bias amino acid choices during Rosetta design.

**Fig S11.**
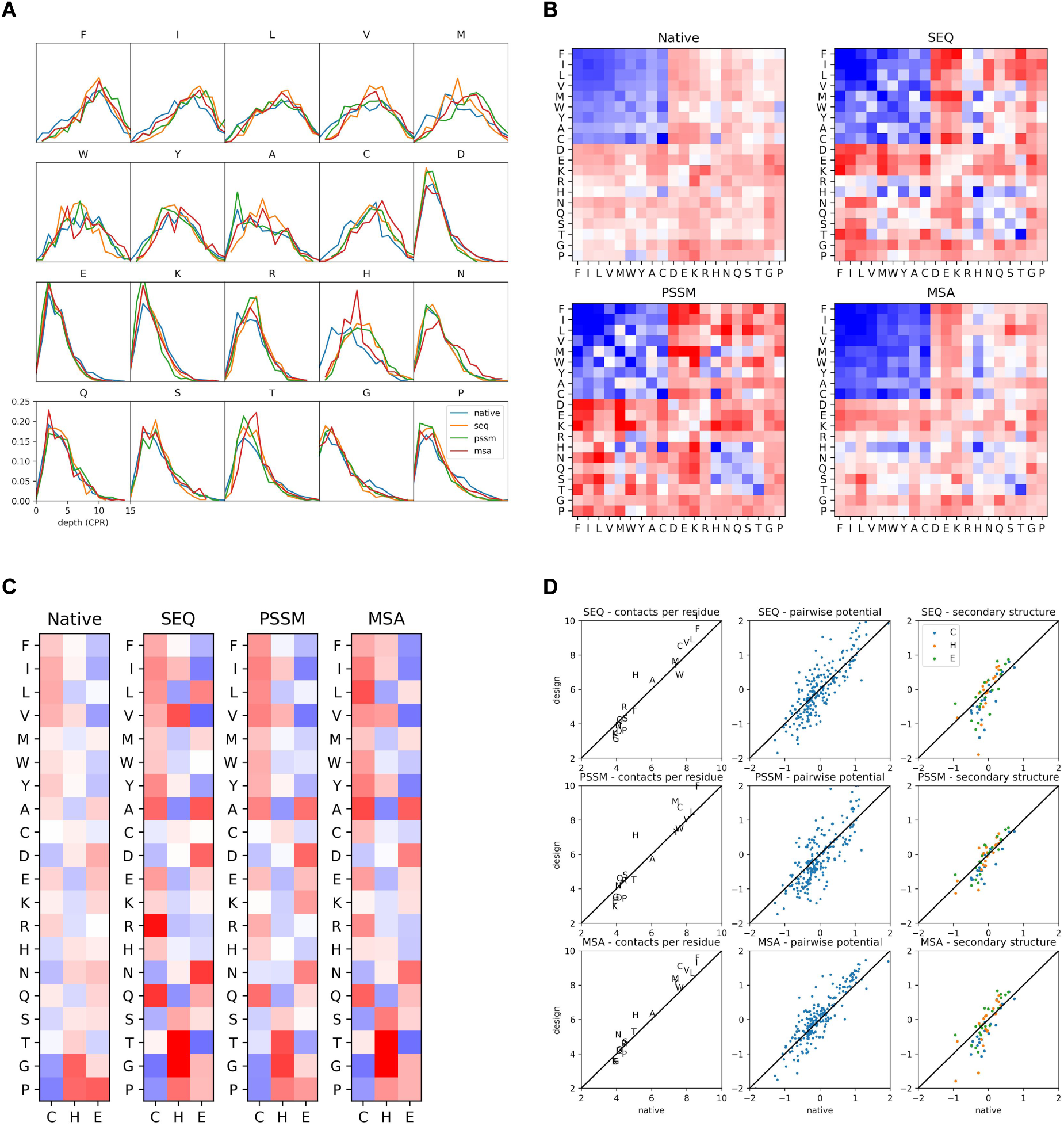
TrRosetta designs recapitulate statistics from the PDB. For the 209 proteins from the dataset described in Supplementary Methods, we designed new sequences with trRosetta using single-sequence, PSSM and MSA mode. (**A**) For each amino acid type, we computed the average number of possible contacts per residue (CPR), using CONFIND (35), with suggested cutoff of 0.01. All three modes of design recapitulate CPR. (**B**) For each design, we threaded the sequence onto the original backbone, and repacked the side-chains using SCWRL4 (36), and defined contact using min(distance) < 5.0. The protocol recapitulates the pairwise potential, defined as *log(p(a,b*|*contact)/(p(a)p(b))*. (**C**) Amino acid propensities for each secondary structure, computed as *log(p(aa*|*ss)/p(aa))*. For (**B-C)**, the scale for the color is −1.5 to 1.5. The results are summarized in **D**.

**Fig S12.**
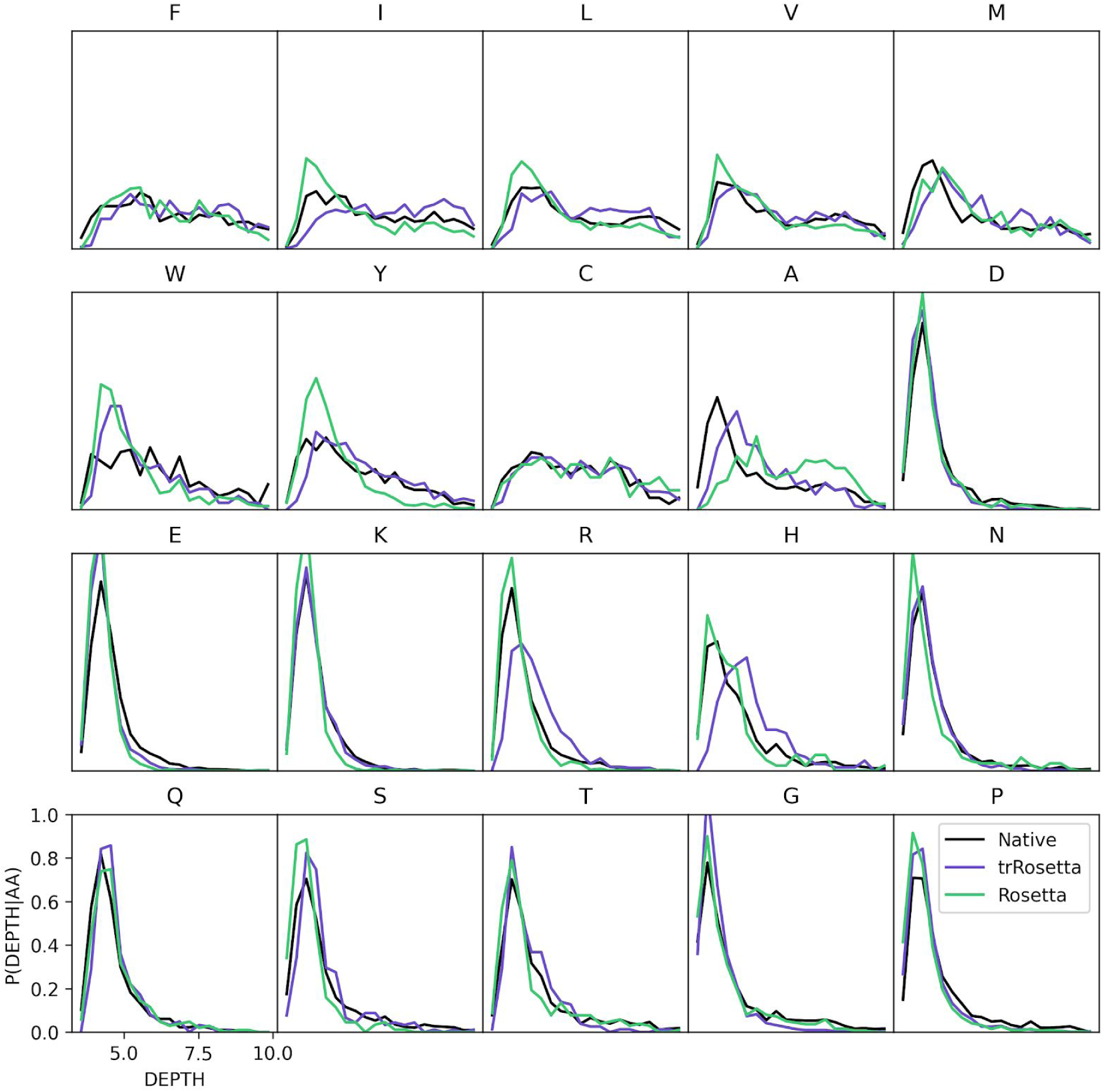
DEPTH distribution for single sequence design using Rosetta and trRosetta on a set of native backbones.

## Supplementary Methods

### Generation of structure-fragment based sequence profiles

To improve the local sequence-structure relationship it can be helpful to restrict and bias sequence design by sequence profiles derived from natural sequences. This has been successfully applied to a range of design problems where native protein families were available (37–40). For *de novo* structures and structure remodeling, where natural sequence alignments are unavailable, fragment-based approaches have been used instead (19, 41). Here, we used the approach from Brunette *et al*. (19), which generates sequence profiles by joining the sequences of matching native structure fragments for each 9-residue-long sliding peptide window in the design. To account for sequence stretches where only a few native fragments are found, we add a small pseudocount to the observed amino acid frequencies (*f* _*a*_):

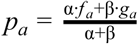

where α is the total number of observed amino acids for that site, β is a constant pseudocount of 50, and *g*_*a*_ is the pseudo frequency computed as:

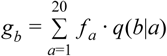

where *q(b*|*a)* is the conditional probability of amino acid *b* given *a* in the BLOSUM62 matrix (42). Finally, we compute half-bit log-odds ratio scores from the pseudocount-corrected frequencies and the background frequencies of each amino acid as in BLOSUM62, *s* = 2*log*_2_(*p*_*a*_/*p*_*a*_, _*background*_), and applied these scores to restrict and bias Rosetta’s FastDesign algorithm (24, 27).

### Generation of trRosetta-based sequence profiles

To make trRosetta-based PSSMs, we first generated 100 sequences for each Foldit-player design following the method described in Fig. S1A, and from those, computed natural log-odds scores by dividing the position-specific amino acid score with the background frequency across all sites.

### Restricting and biasing Rosetta sequence design

Sequence design with Rosetta was performed with FastDesign (24, 27), using the Rosetta all-atom energy function (28) with beta16_nostab weights. We applied three different methods for restricting and biasing amino acid choices during FastDesign. In the first method (LayerDesign, (18)), amino acid identities were restricted based on the site’s surface exposure, secondary structure type and helix capping, generally disallowing hydrophobic amino acid on the surface, and hydrophobic amino acids in the core. For the two other methods (fragment-based PSSMs and trRosetta-based PSSMS), we disallowed amino acids with PSSM scores less than −1, and biased the energy function by the −0.3 * PSSM score.

### TrRosetta amino acid bias weight optimization for position probability matrix design

The composition-based loss proposed in Eq. 3 biases the composition of each designed protein towards the mean composition of proteins in the PDB. This is not always desirable, as the ideal composition for any specific protein could deviate from the mean composition of the PDB. Instead, to correct for the intrinsic amino acid biases in trRosetta, we implemented a new loss to maximize the agreement between position-specific amino acid frequencies (sequence profile) of natural proteins and those designed by trRosetta. Specifically, the new loss biases the frequency of each possible amino acid at each position in a designed sequence profile via a fitted weight:

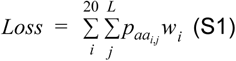

Where 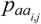 is the probability of the *i* ‘th canonical amino acid at residue position *j*, and *w*^*i*^ is the bias loss weight associated with amino acid *i*. Eq. S1 takes a form that must be enforced at the sequence profile level (Fig. S1B, softmax-PSSM), as we found this performed better than applying the loss to the sequence sampled from the profile. As such, we analogously implement the composition-based loss seen in Eq. 3 at the sequence profile level to maintain direct comparability during the following analysis.

To fit the weights *w*^*i*^ we used Nelder-Mead simplex optimization (43), minimizing the total KL-divergence between a set of sequence profiles derived from native sequence alignments and profiles generated by outputting the softmax-PSSM from trRosetta PSSM design mode (Fig. S1B):

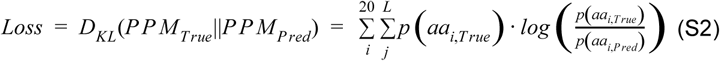

To build a training set for the optimization, we collected native structures and their corresponding sequence alignments from Hiranuma *et al*. (44), and reduced the redundancy of the set so that no structure had more than 30% sequence identity to any protein in the trRosetta training dataset. This resulted in 209 training proteins, whose sequence profiles were then computed using PSI-BLAST (45, 46).

We ran the minimization process for approximately 3 days using 20 rtx2080 GPUs in parallel. Relative to using the original formulation of Eq. 3, this improved the profile of designed sequences, decreasing the per-residue KL-divergence from 0.75 to 0.71 (5.1%).

